# Microbiome complexity shapes host transcriptional networks across the ileum-liver axis

**DOI:** 10.64898/2026.11.16.767119

**Authors:** Amit Coomar Halkhoree, Alex Grannell, Claire Pearson, Elizete Araujo, Rebecca Jeffery, Kelsey Jones, Gregor Rot, Eric Galvez, Fiona Powrie, Petar Scepanovic, Nicholas Ilott

## Abstract

The microbiome plays an increasingly recognised role in development, metabolism and immunity across all segments of the small and large intestine, as well as more distant organs such as the liver. In the terminal ileum – a key site of nutrient absorption – the presence of microbes dictates the development of secondary lymphoid structures (Peyer’s patches), mediates bile acid metabolism, and impacts absorption. In part, coordinated regulatory networks in the gut impact liver function and immunity through gut-liver connectivity and the movement of immune cells and bioactive small molecules that includes bile acids and other bacterial metabolites.

We aimed to examine the extent of microbiome involvement in ileal transcription, where subtle changes in co-expression networks can be tuned by microbiome diversity, composition and function. We surveyed ileal transcriptomes across mouse models which vary in microbiome complexity ranging from germ-free (GF, n = 15), the Oligo-Mouse-Microbiota-12 (MM12, n = 15), and specific pathogen free (SPF, n = 15).

**IMPORTANCE:** We observed that the ileal MM12 community was taxonomically similar to SPF ileal communities, dominated by *Lactobacillus* species. However, metagenome-scale metabolic modelling (MSMM) revealed a more restricted predicted metabolic repertoire in this community which could impact host functions. In ileal transcriptomes, we corroborate previously identified alterations in GF relative to SPF mice and crucially show that mice harbouring the MM12 community have transcriptional profiles which are more similar to GF mice than SPF mice, where recapitulation of gene expression to levels observed in SPF mice varies by gene and biological process. Microbiome complexity alters ileal gene co-expression network topology, where the low complexity MM12 community restores coupling of fatty acid metabolic processes observed in SPF mice. We further identify *Faecalibacterium duncaniae* as predicted to be a key species associated with multiple and varied host functions that includes fatty acid metabolism and innate immune networks. Finally, liver transcriptomes in GF and minimally diverse microbiomes reflect altered ileal processes in terms of responses to lipid absorption and circulation.

## INTRODUCTION

The ileum and liver are interconnected through a complex, yet not fully understood, communication network that plays a crucial role in maintaining metabolic and immune homeostasis.^1^ They are linked via the bile ducts, portal vein, lymphatics and systemic circulation, with their interactions being essential for the efficient breakdown and absorption of dietary components, as well as enterohepatic circulation of bile acids.^2^ The ability of gut microbes to produce bioactive compounds, to transform bile acids, and to act as an antigenic pool therefore supports a role for the gut microbiome in mediating ileal and liver immunity and physiology.

Mice that are born and raised under sterile conditions (germ-free, GF) have overt physiological differences compared to conventionally raised, specific pathogen free (SPF) laboratory mice. These are reflected in lower body weight, enlarged caeca and reduced adiposity. These deficits result from an inherent lack of microbe-dependent carbohydrate fermentation, nutrient metabolism and bile acid modifying processes.^3^ Broad impacts of microbiome presence on transcriptional programs across a variety of tissues have been described using single-cell and spatial transcriptomic technologies,^4–8^ providing key insights into microbially-mediated immune and metabolic functions in mice. These surveys have revealed that presence of a microbiome determines the development, differentiation and activation of lymphocytes including IgA-producing plasma cells and effector T cell subsets such as Th17 and Treg, as well as antigen processing and presentation processes in myeloid populations in the ileum.^9^ Further, microbial coupling of lipid metabolism across the ileum and liver has been revealed through coordinated alterations in gene expression of lipid absorption, transport and handling pathways between ileal epithelial cells and hepatocytes.^4^

The specific microbes, enzymatic reactions, and signalling components that mediate aspects of these pathways are beginning to emerge. For example, colonisation of a single strain of bacteria, segmented filamentous bacteria (SFB), can recover the reduced development and activation of Peyer’s patches observed in GF mice^10^, as well as direct CD4 T cells towards a Th17 phenotype in SPF mice.^11,12^ Furthermore, microbiome-mediated regulation of bile acid homeostasis impacts ileal epithelial cells to control barrier integrity^13^ and lipid metabolic processes.^14^ Indeed, in an intricate network of tuneable cross-kingdom interactions, presence of a microbiome increases biliary taurine-conjugated bile acid concentrations, leading to increased conversion of phosphatidylcholine to lysophosphatidylcholine and reduced lipid absorption in the small intestine.^14^ In addition to metabolic impacts, the ileal microbiome imparts long-range effects on liver immunity^15^ that together underscore the importance of understanding how specific microbes and microbial consortia impact cross-tissue functional networks.

There is strain-dependent variation in how microbes in the ileum influence host metabolism and immunity, yet how microbiome diversity, microbiome composition, and microbiome function impact on the host are not completely clear. Low complexity gnotobiotic models - animals with a defined microbial consortium - help to reduce the complexity of microbe-microbe and microbe-host interactions observed in SPF mice, whilst retaining broad microbiome functional capacity. One example is the Oligo-Mouse-Microbiota 12 (hereafter referred to as MM12) community, a synthetic 12-member consortium designed to recapitulate the 5 major phyla found in the mouse colon, that recapitulates broad functions of SPF mice that includes colonisation resistance^16^ colonic immune profiles^17^ and colonic disease phenotypes.^18^ However, while this has become an established model of colonic microbiome functionality, less is known about how minimal microbiome communities such as the MM12 affect functions of the ileum and downstream impacts on the liver. Nevertheless, it is clear that diversity *per se* is not the sole mediator of host metabolism, as MM12 mice have higher fat mass than SPF and GF mice and their respiratory exchange ratio (RER) recapitulates SPF or GF RER in a circadian-dependent manner.^19^ In addition, both liver and plasma metabolite profiles in MM12 mice are more similar to GF than SPF mice,^19^ suggesting that there are microbiome functions that are lacking in the MM12 which are important for liver metabolic processes.

In this study, we expand on a growing literature that describes the impact of microbiome complexity on host functions. Looking specifically at the terminal ileum, we uncover selective transcriptional networks which are mediated by microbiome diversity. We demonstrate that a minimally complex microbiome (MM12) can restore coupling of fatty acid metabolic networks that are lost in GF mice yet is unable to restore gene co-expression networks related to muscle cell differentiation. By analysing correlations with species present in the SPF community, we identify *Faecalibacterium duncaniae* as a potential mediator of multiple and varied host functions that includes metabolic control of immune-related transcription.

## RESULTS

### Microbiome complexity shapes predicted community metabolic output and niche occupancy

The MM12 consortium was designed to recapitulate major functions of the colonic microbiome. However, colonisation of this consortium and its functional capacity in the small intestine have not yet been assessed.

We first wanted to understand how the minimally diverse MM12 community relates to an SPF community in terms of diversity and functional capacity in the terminal ileum. We took ileal contents from C57BL/6 mice which had been colonised from birth with either the MM12 community or SPF community and assessed the relative abundance of taxa using either qPCR (MM12) or shotgun metagenomic sequencing (SPF). The ileal SPF community was significantly more diverse than the MM12 community (**Fig. 1a**) and differed globally in terms of species relative abundance (**Fig. 1b**). At higher taxonomic ranks, the MM12 recapitulates SPF ileal microbiomes, with major representation of phylum Bacillota (formerly Firmicutes) across samples (**Fig. 1c**). Likewise, at family and species levels, similarities between these two communities are evident, with high representation of family Lactobacillaceae and *Lactobacillus spp.* in both MM12 and SPF samples (**Fig. 1c**).

**Fig. 1.**
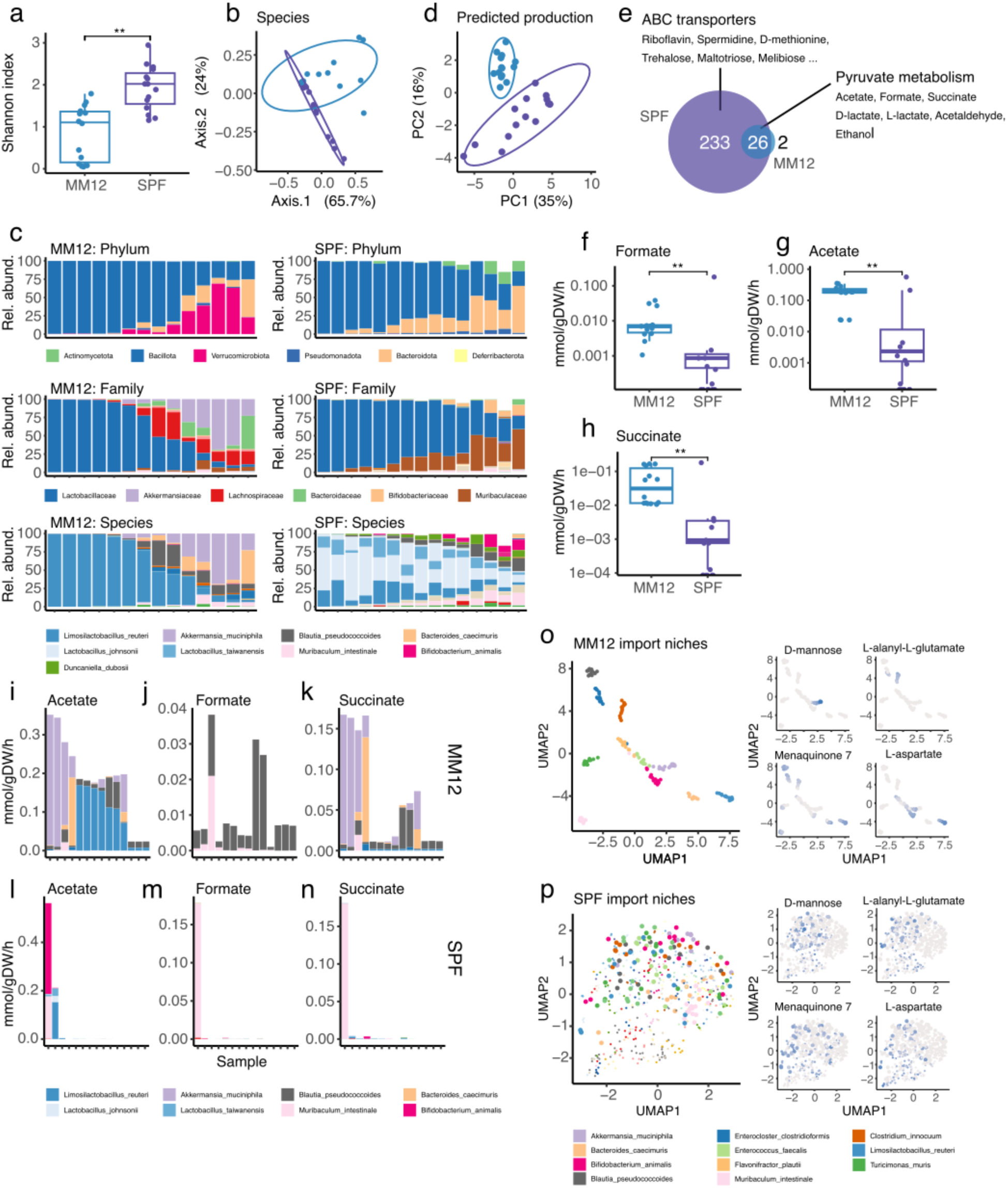
The minimally diverse MM12 community has reduced predicted metabolic diversity than SPF. (a) Comparison of Shannon diversity index between ileal MM12 (n = 14) and SPF (n = 15) mice. Results based on Wilcoxon rank sum test. (b) PCoA based on Bray-Curtis distance. Points are coloured by microbiome status. (c) Stacked barcharts displaying the percentage of reads mapping to different phyla, families and species across MM12 and SPF mice. (d) PCA plot of predicted production capacity of MM12 and SPF communities. (e) Venn diagram of metabolites predicted to be produced by the MM12 and SPF communities in the ileum. Changes in predicted flux of products of pyruvate metabolism (f) formate, (g) acetate and (h) succinate between MM12 and SPF ileal communities. Species contributions to export fluxes of (i) acetate in MM12, (j) formate in MM12, (k) succinate in MM12, (l) acetate in SPF, (m) formate in SPF and (n) succinate in SPF). (o) UMAP of predicted import fluxes of each species present in the MM12 across samples. Examples of metabolites that distinguish species import niches are shown (p) UMAP of predicted import fluxes of each species present in SPF across samples. Examples of metabolites that distinguish MM12 import niches are shown. Species that belong to the MM12 community are in large point and non-MM12 members in smaller point. UMAPs were generated using default configuration in the umap v0.2.10.0 package in R.

Nevertheless, key differences exist. Verrucomicrobiota are enriched in a subset of MM12 samples, where Bacteroidiota are enriched in a subset of SPF samples (**Fig. 1c**). Enrichment of Verrucomicrobiota phylum in MM12 is reflected at lower taxonomic ranks with Akkermansiaceae and *Akkermansia muciniphila* being highly represented at the family and species levels, respectively (**Fig. 1c**). Family Muribaculaceae is present in both SPF and MM12 communities yet reaches higher relative abundance in a subset of SPF samples relative to MM12 (**Fig. 1c**).

To understand the potential functional consequences of diversity and/or species compositional differences in MM12 relative to SPF communities, we performed metagenome-scale metabolic modelling (MSMM) using MICOM.^20^ Using a bespoke dietary input (see Methods) and relative abundance, we predicted sample-wise metabolic flux and interpreted both the production capacity of the community, as well as exchange fluxes that account for cross-feeding. Using total metabolite production capacity (i.e. the sum of species-level metabolite exports), we could demonstrate that global community metabolism varies in MM12 mice relative to SPF mice (**Fig. 1d**). The differences are, in part, related to a more restricted metabolic production capacity in the MM12, with 28 metabolites being predicted to be exported from the MM12 community, compared to 259 from the SPF community (**Fig. 1e**). Of note, the MM12 retain core carbohydrate metabolic capacities, where pyruvate metabolism which results in the production of acetate, formate, succinate, D-lactate and L-lactate, is a significantly enriched pathway in metabolites shared between MM12 and SPF communities (**Fig. 1e**). In contrast, metabolites which constitute substrates such as riboflavin and spermidine and are exported/imported through ABC transporters are enriched in metabolites that are predicted to be produced to appreciable levels from the SPF community only (**Fig. 1e**).

Although the MM12 are predicted to have a somewhat constrained metabolic capacity relative to SPF, overall production of metabolites from carbohydrate metabolism is predicted to be higher, which includes increased predicted production of formate, acetate and succinate from MM12 relative to SPF (**Fig. 1f-h**). These increases are reflected in increased relative abundance of species which are predicted to be the largest exporters of these metabolites including *Akkermansia muciniphila* for acetate and succinate production, and *Blautia (pseudo)coccoides* for formate production (**Fig. 1i-n**).

We next wanted to assess to what degree an increased diversity in the SPF community dictates species niche occupancy – a feature that could explain differential metabolic capacity. To visualise metabolic niches, we took predicted species-level import fluxes and visualised clustering of species across samples using Uniform Manifold Approximation and Projection (UMAP). From these niche maps, we observed that members of the MM12 clustered into distinct import niches that were similar across samples (**Fig. 1o**). In contrast, SPF import niches were much broader and overlapping between species with less similarity across samples (**Fig. 1p**). More varied import niches for MM12 members that are part of the SPF community were also observed, suggesting an impact of the surrounding community on metabolite availability for import (**Fig. 1p**). Predicted imported metabolites were spread across more species in general in the SPF community relative to MM12 community (**Supplemental Material Fig. S1**), supporting a model of nutrient competition in more diverse communities (examples provided in **Fig. 1o-p**).

Together, our data suggest that the ileal MM12 broadly recapitulates the more diverse SPF community in terms of taxonomy, capturing the dominance of Bacillota and particularly Lactobacillaceae. However, the predicted metabolic production capacity of the MM12 community is more constrained and supports separation of species into metabolic niches that are more well defined than those of members in the SPF community.

### Microbiome complexity directs ileal transcriptional profiles

Given the observed differences in taxonomy and predicted metabolism in the lower diversity MM12 community relative to the SPF community, we next sought to understand the impact of community differences on transcriptional profiles in the ileum.

Given previous literature on changes to ileal transcriptomes in GF mice relative to SPF mice,^4,6^ we first wanted to test the extent to which we could verify previous observations in our dataset. As expected, clustering of samples using PCA demonstrated broad transcriptional changes in GF relative to SPF mice in the ileum (**Supplementary Material Fig. S2a**, Adonis test expression (vst) ∼ microbiome status, *R*^2^ = 0.14, p = 0.001). Using DESeq2, we found a total of 2210 genes as differentially expressed (padj < 0.05) between GF and SPF mice (**Supplemental Table 1**). We confirmed previous observations of reduced expression of genes related to adaptive immune responses and increased expression of genes related to lipid metabolic processes in GF mice relative to SPF mice (**Supplementary Material Fig. S2b** and **Supplemental Table 2**). Genes involved in plasma cell function and soluble antibody production including *Igha* and *Ighb2b* (**Supplementary Material Fig. S2c**) were reduced in GF mice, as well as genes involved in lymphocyte development and survival, including *Il7* and *Il7r* (**Supplementary Material Fig. S2c**), as expected. We were also able to confirm the increased expression of lipid handing and transport genes (*Apoa1, Apoa4*, *Fabp1* and *Fabp2*, **Supplementary Material Fig. S2c**) in GF ileum relative to SPF mice, that likely relate to zonation differences and thickening of epithelial barrier when the microbiome is not present.^4^

Next, we assessed the ability of a minimal microbiome community (MM12) to restore gene expression to levels observed in SPF mice. Although MM12 mice recapitulate broad functionality of a more complex colonic microbiome, transcriptional profiles of MM12 mice in the ileum were more similar to GF profiles than SPF profiles (**Fig. 2a**). By comparing differential expression results, we were able to categorise GF-associated genes as being either “recovered” (non-significant difference in MM12 vs. SPF) or “not recovered” (significantly different in MM12 vs. SPF) through colonisation with the MM12 community (**Supplemental Table 3**). We found that most genes that are altered in GF conditions are not recovered through colonisation with the MM12 community (**Fig. 2b**). Visualisation of microbiome-mediated genes in a heatmap revealed that genes classified as recovered in MM12 mice were highly variable in expression between individual MM12 mice and largely were somewhat intermediate between SPF and GF (**Fig. 2c**). Pathways associated with recovered gene expression in MM12 mice were those involved in cell activation and RNA processing (**Fig. 2d**), where genes that were not recovered were associated with immune response and lipid metabolism (**Fig. 2d**).

**Fig. 2.**
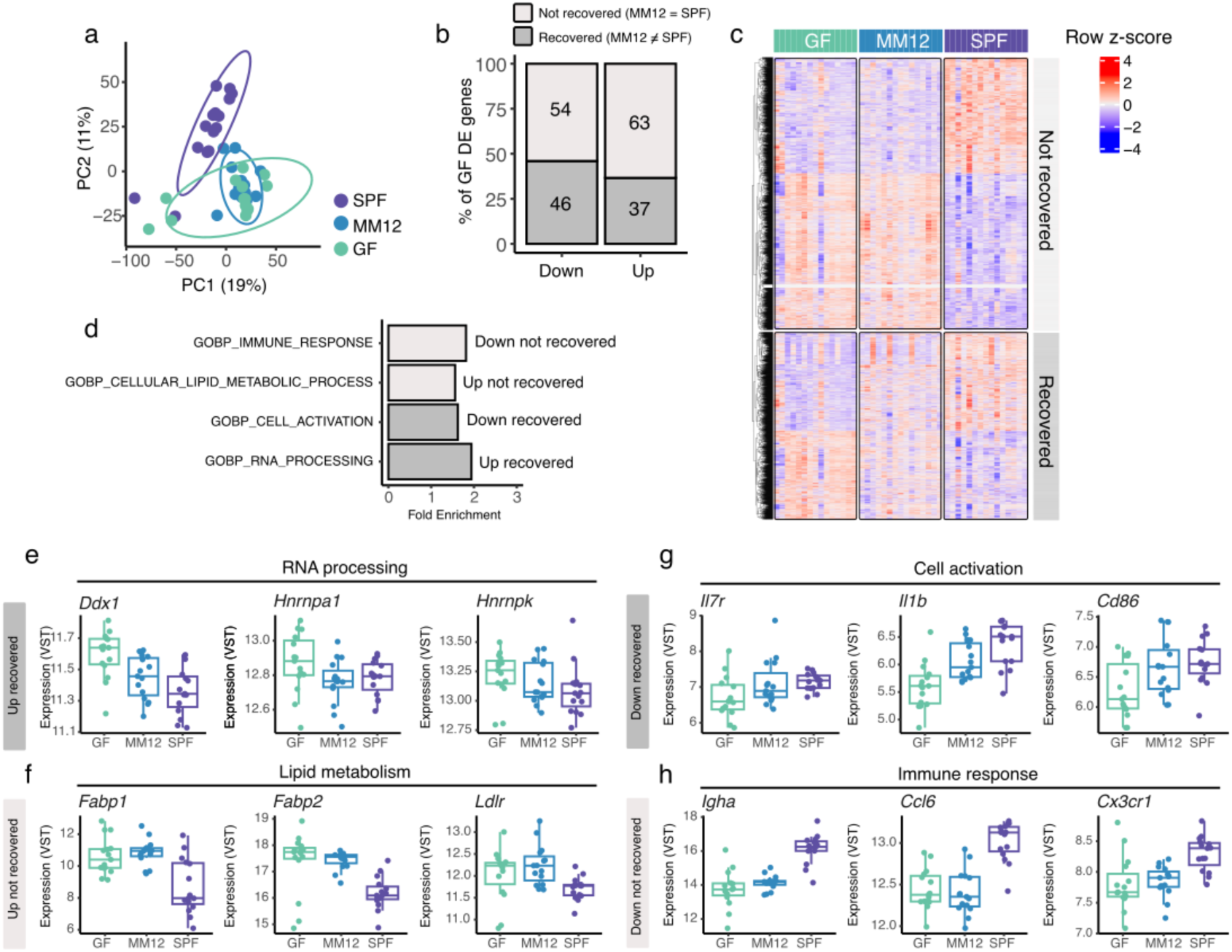
**Transcriptomic profiling of the ileum across microbiome states**. (a) Principal components analysis (PCA) and two-dimensional ordination of profiles across SPF, MM12 and GF ileum samples (n = 15/group). (b) The proportion of genes recovered and not recovered to SPF levels in MM12 mice in the ileum. (c) Visualisation of microbiome-associated genes (i.e. significantly different in GF vs. SPF mice, padj < 0.05) and their MM12 recovery status in the ileum. (d) Pathway (gene ontology) over-representation analysis of genes dependent on MM12 recovery status in the ileum. (e-h) Boxplot visualisations of selected genes based on their MM12 recovery status and GO pathway association/functions.

As noted previously, the level of recapitulation of MM12 gene expression to SPF levels varied by gene (**Fig. 2e-h**). We observed that the minimally diverse MM12 community could suppress the expression of splicing factors (**Fig. 2e**, *Ddx1*, *Hnrnpa1, Hnrnpk*) relative to GF mice. However, genes involved in lipid metabolism that are increased in the ileum of GF mice (*Fabp1*, *Fabp2*, *Ldlr*) were not suppressed by the MM12 community to levels observed in SPF mice (**Fig. 2f**). Further, we observed varying patterns of recovery in immune responses. For example, genes involved in activation and facilitating the development and survival of memory T cells (*Il7r*, *Il1b* and *Cd86*) were increased in MM12 ileal samples to similar levels observed in SPF mice (**Fig. 2g**), supporting a potential induction of bacterial antigen presentation and commensal T cell memory. However, other immune responses that are typical of plasma cells (*Igha* expression) and myeloid cell recruitment (*Ccl6*) and function (*Cx3cr1*) remained significantly reduced in MM12 mice relative to SPF mice, and similar to levels observed in GF (**Fig. 2h**).

Together, our data suggest that the presence of a microbiome alone is not sufficient to recover transcriptional profiles observed in SPF mice, such as lipid handling and some adaptive immune responses. Furthermore, missing features in the MM12 community impact transcription in the ileum, particularly in lipid metabolism and immune response.

### The low diversity MM12 community exerts specific impacts on the ileal transcriptome

In addition to recovery of transcriptomic programmes from GF levels to SPF levels, we were interested to understand if the MM12 community had any specific associated changes. We performed differential expression analysis comparing MM12 to both SPF and GF mice and identified those genes that were increased or decreased in both comparisons. We identified 209 up and 218 down-regulated genes in MM12 mice relative to both SPF and GF (**Supplemental Table 3** and **Supplemental Table 4**). Using overrepresentation analysis we found 8 GO BP significantly enriched among genes upregulated and 157 among downregulated genes (**Supplemental Table 5**). We observed coherent regulatory pathways in up and downregulated genes, where genes involved in transcriptional responses to cellular stress (*Ppp1r15a, Hif1a, Ddit3* and *Atf3*) were upregulated (**Fig. 3a**), and genes involved in DNA replication and cell proliferation (*Mki67, Top2a, Cdc20, Cdca2*) were downregulated (**Fig. 3b**).

**Fig. 3.**
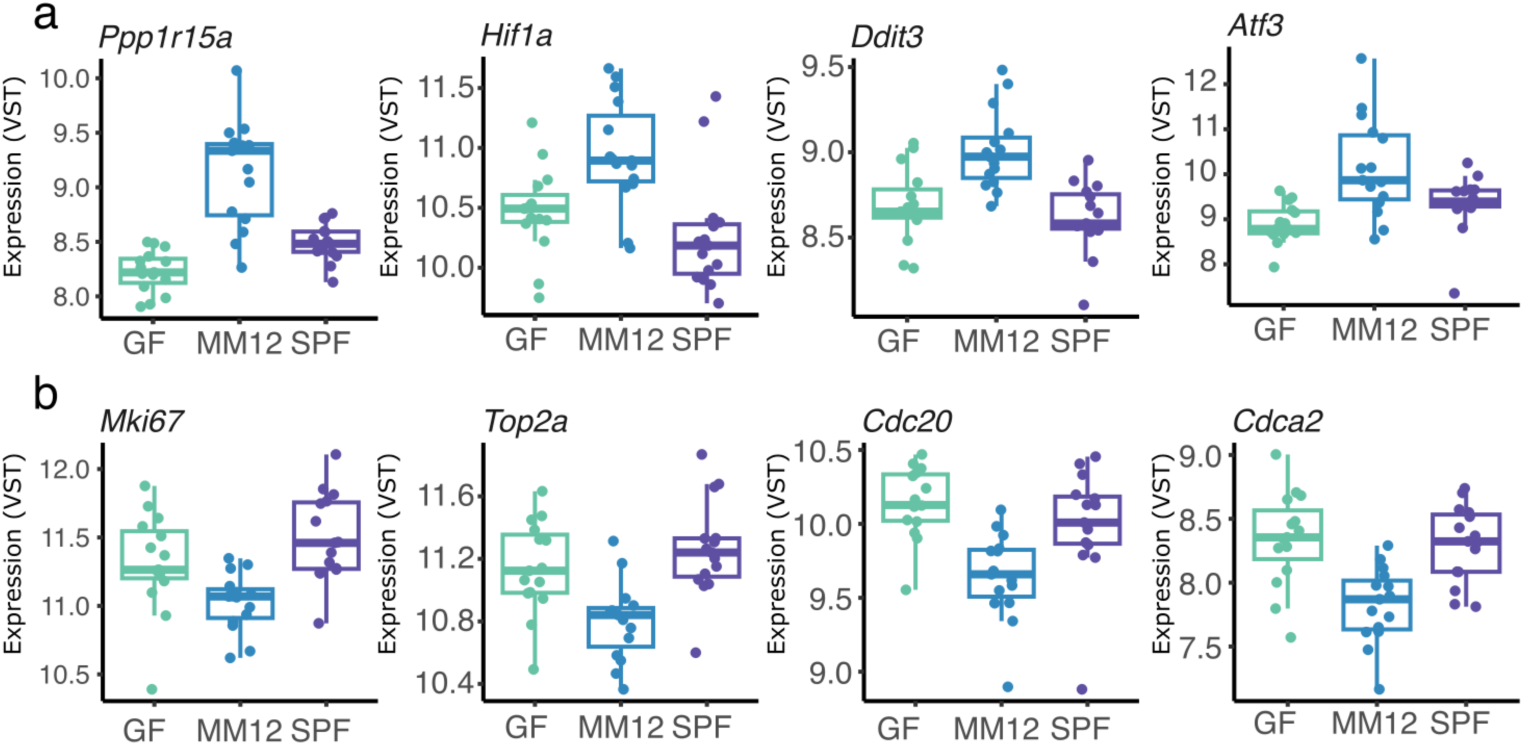
**MM12 mice have distinct transcriptional programmes relative to GF and SPF mice**. (a) Expression (VST) levels of a selection of genes that are annotated as genes involved in the GO BP GOBP_REGULATION_OF_DNA_TEMPLATED_TRANSCRIPTION_IN_RESPONSE_TO_STRESS pathway. (b) Expression (VST) levels of a selection of genes that are annotated as genes involved in the GO BP GOBP_CELL_CYCLE_PROCESS pathway.

These data support transcriptional mediation by a minimal microbiome which is distinct from that of a more complex community. This supports a concept of transcriptional control above the mere presence or absence of an ileal microbiome community.

### A minimal microbiome restores metabolism-related ileal co-expression networks

Absolute differences in gene expression provide insights into up- and down-regulated processes or changes in cell composition across microbiome states. However, we also designed our experiment with sufficient samples per group to assess gene co-expression networks and how microbiome status impacts gene coupling and connectivity.

First, we generated a baseline ileal gene co-expression network and subsequent co-expression modules using Weighted Gene Co-expression Network Analysis (WGCNA) using data from SPF mice only. We identified 19 modules which contained between 120 (ILE18, steelblue) and 2916 (ILE6 - blue) genes. Each module was assigned functional annotations based on GO BP overrepresentation analysis. We found significant pathway enrichment for all ileum modules (**Supplemental Table 6**), supporting functional coherence in our co-expression network. Coordinated regulation of genes in each module was confirmed using PCA, where the range of variance explained by PC1 for each module (**Fig. 4a**) was between 41% (ILE6, blue module, proteolysis) and 70% (ILE11, green, hormone/fatty acid metabolic process).

**Fig. 4.**
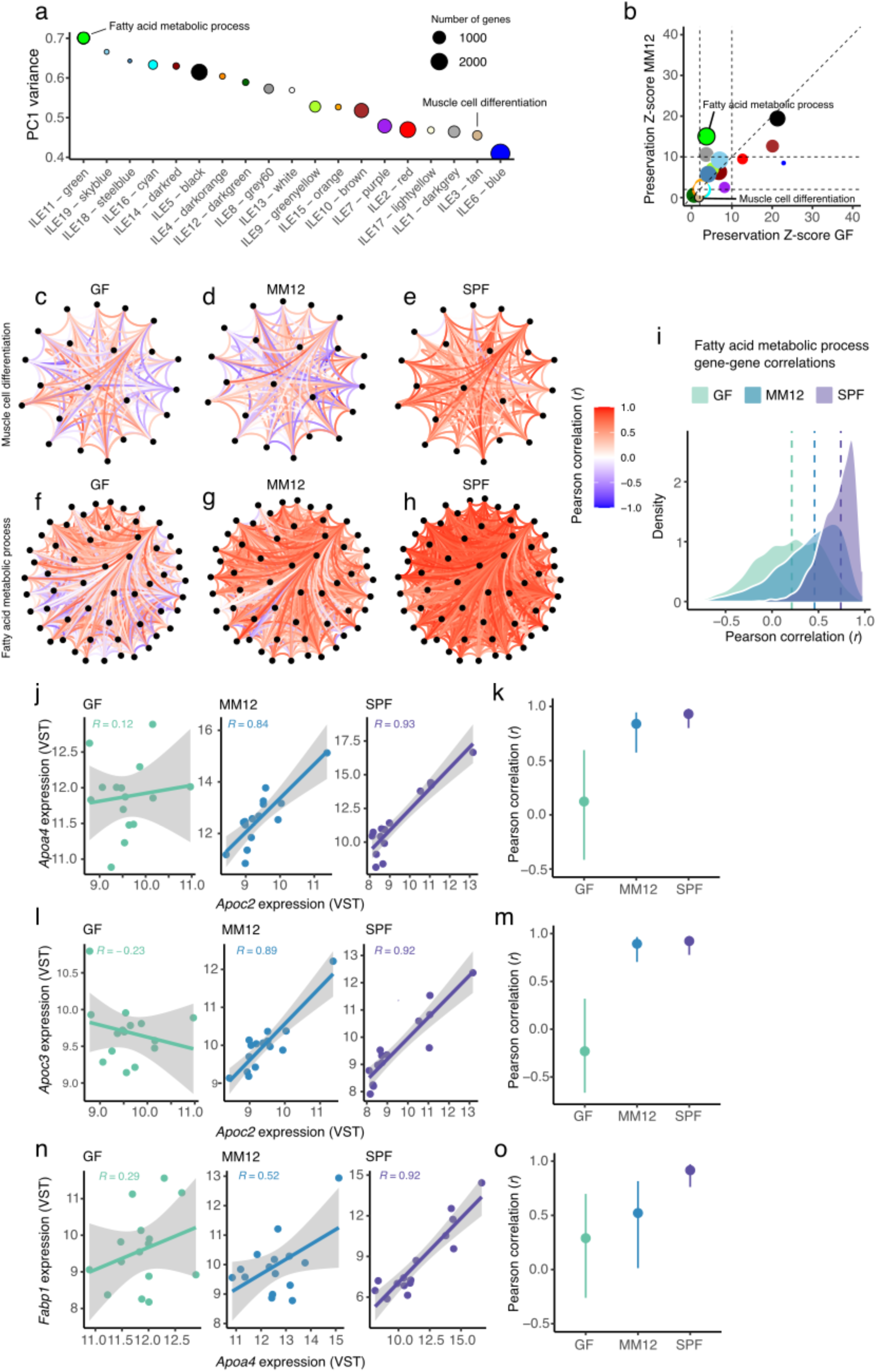
Microbiome complexity shapes gene co-expression networks in the ileum. (a) Modules defined by WGCNA in SPF ileum samples (n = 15). Colours represent the assignment by WGCNA. Size of points is determined by the gene set size for each module. (b) Preservation Z Summary scores for GF and MM12 samples based on results from the WGCNA function modulePreservation(). Genes contributing to the enrichment signal for the muscle cell differentiation GO BP and their correlations are shown as graphs for (c) GF (d) MM12 and(e) SPF samples. Genes contributing to the enrichment signal for the fatty acid metabolic process GO BP and their correlations are shown as graphs for (f) GF (g) MM12 and (h) SPF samples. (i) Overall connectivity distribution (Pearson correlations) for genes contributing to the fatty acid metabolic process GO BP across groups. (j) Comparing correlations between GF, MM12 and SPF mice for *Apoc2* and *Apoa4*. (k) Correlation estimate and 95% confidence intervals for correlations between *Apoc2* and *Apoa4* across groups. (l) Comparing correlations between GF, MM12 and SPF mice for *Apoc2* and *Apoc3*. (m) Correlation estimate and 95% confidence intervals for correlations between *Apoc2* and *Apoc3* across groups. (n) Comparing correlations between GF, MM12 and SPF mice for *Fabp1* and *Apoa4*. (m) Correlation estimate and 95% confidence intervals for correlations between *Fabp1* and *Apoa4* across groups.

To understand the impact of microbiome complexity on co-expression networks, we used WGCNA module preservation analysis.^21^ This allowed us to ask whether each SPF-defined module was preserved in GF and MM12 mice, and to what extent. We found that the majority of SPF modules were moderately and similarly preserved in GF and MM12 ileum (**Fig. 4b**, preservation Z-Summary scores > 2) although there were exceptions. For example, ILE3 (tan) module which was enriched for genes involved in tissue organisation (GO BP terms include blood coagulation and muscle cell differentiation, see **Supplementary Table 6**) was not well preserved in neither GF nor MM12 ileum. This was visually confirmed by looking at gene expression correlation between genes that contributed to the enrichment of the GO BP term muscle cell differentiation (n = 23 genes, **Supplementary Table 6**) for this module across microbiome states (**Fig. 4c-e**). In contrast, there was a clear difference in preservation between MM12 and GF mice for ILE11 (green module) which is associated with genes involved in fatty acid metabolism (**Fig. 4b**). This module was strongly preserved in MM12 ileum but weakly preserved in GF ileum (**Fig. 4b**) suggesting that the presence of a microbiome couples genes involved in fatty acid metabolic processes. Taking genes in the GO BP pathway fatty acid metabolic process, we visually observed the increased connectivity in MM12 mice relative to GF mice (**Fig. 4f-i**). Furthermore, we could statistically demonstrate that correlations between key genes involved in assembling lipids into chylomicrons (*Apoa4, Apoc2, Apoc3*) are reduced in GF mice relative to both SPF and MM12 mice (**Fig. 4j-m**), supporting a role for the microbiome in coupling lipid assembly processes. In addition, we observed that the full repertoire of connections were not preserved in MM12 mice for this module, where correlations between lipid binding protein, *Fabp1* and the apolipoprotein, *Apoa4* were intermediate for MM12 mice (**Fig. 4n-o**).

Taken together, these data support complex and varied impacts of microbiome complexity on coupling host transcriptional processes in the ileum.

### *Faecalibacterium duncaniae* is a key species in the ileum that is associated with diverse lipid-handling associated transcriptional programmes

To determine associations between species and host transcriptional programmes, we correlated metagenomics-derived species relative abundances with module eigengenes in SPF ileal data (n = 46 species, n = 19 modules). We identified 14 strong (|Spearman’s rho| > 0.6) species-module correlations with 4/14 (28%) containing *Faecalibacterium duncaniae* (**Fig. 5a**). Placing *F. duncaniae* in a network with all host modules (**Fig. 5b**), we visualised its positive associations with modules annotated as those involved in fatty acid metabolism (**Fig. 5c**, green module) and RNA processing (**Fig. 5d**, including chromatin organisation and splicing, purple module). *F. duncaniae* was negatively associated with modules annotated as containing genes involved in innate immune responses (**Fig. 5e**, brown and **Fig. 5f**, black modules). Through exploring *F. duncaniae*-gene correlations within these associated modules, we observed that some of the strongest positive associations were between *F. duncaniae* and genes within these modules that are involved in metabolic processes. This included the previously described genes involved in chylomicron assembly and lipid metabolism (**Fig. 5g-j**) as well as *Mgam* (**Fig. 5k**), which encodes the brush-border maltase-glucoamylase that is key for breaking down starch into absorbable glucose.^22^

**Fig. 5.**
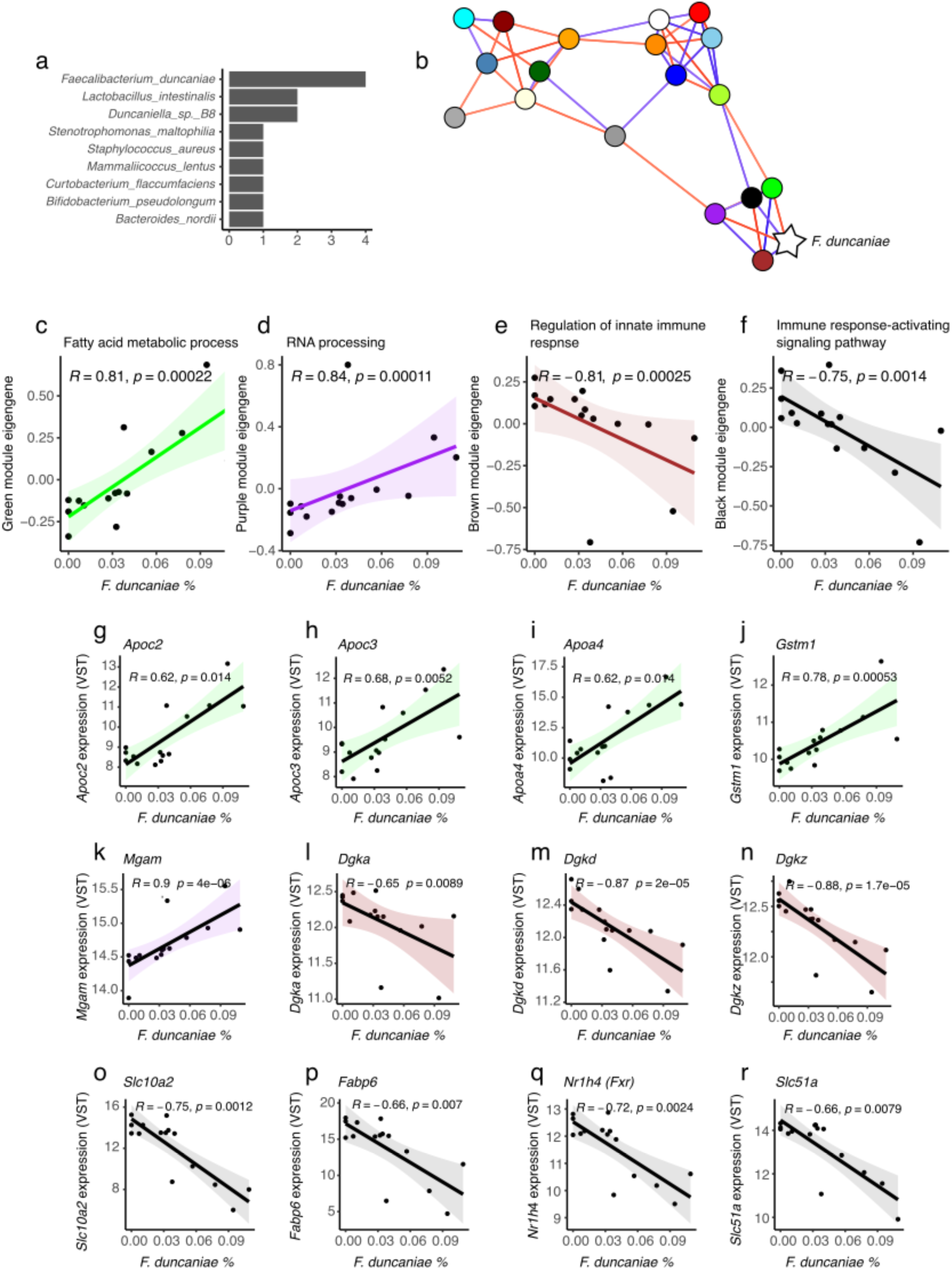
***F. duncaniae* is associated with ileal transcription programmes**. (a) Species and the number of strong (|Spearman’s Rho| > 0.6) correlations observed (b) Correlation network based on Pearson’s correlation between module eigengenes and including *F. duncaniae*. Colours indicate module colours from WGCNA. Scatterplot visualisations of correlations between *F. duncaniae* and (c) ILE-11 green, (d) ILE-7 purple, (e) ILE-10 brown and ILE-5 black module eigengenes. Scatterplots of genes of interest expression levels from each of the associated modules is shown in (g-r) and includes those involved in fatty acid metabolic processes, starch breakdown, diacylglycerol conversion, and bile acid homeostasis.

While modules that were negatively associated with *F. duncaniae* relative abundance were annotated as innate immune response modules, we observed that the strongest correlates of *F. duncaniae* were upstream metabolic modulators of innate defence responses rather than canonical downstream effectors. For example, the strongest correlates in the brown module (regulation of innate immune response) were the diacylglycerol kinases *Dgka, Dgkd* and *Dgkz*. These encode enzymes which convert cellular diacylglycerol to phosphatidic acid to promote pro-inflammatory cytokine production from cells such as dendritic cells and macrophages, ^23^ markers of which (e.g. *Alpk1*, *Tifa*, *Tlr2*, *Tlr5*) are present in this module (**Supplementary Table 7**). Likewise, the black module (immune response-activating signaling pathway) contains multiple genes involved in innate cell sensing and signaling (e.g. *Tlr4*, *S100a14*, *Ikbkg*, *Tnf*, **Supplementary Table 6** and **Supplemental Table 7**), yet the strongest associations with *F. duncaniae* are with genes involved in bile acid uptake (*Slc10a2*), signalling (*Fabp6*, *Nr1h4*) and export (*Slc51a*) (**Fig.5o-r**). These associations likely reflect interactions between the epithelium and innate immune cells where, for example, bile acid availability may impact on activation of macrophages through Fxr or alternative bile acid signalling pathways.^24^

Overall, these data support an association of *F.duncaniae* with multiple and varied host metabolic and immune functions that have bile acid homeostasis and lipid metabolic processes at their core.

### Microbiome complexity mediates long-range systemic interactions

Given our observations of some key changes to lipid metabolic transcriptional programmes in GF and to some extent MM12 mice, we were interested to understand if these were associated with effects in the liver. Lipid assembly in the small intestine is heavily linked to liver responses, as chylomicrons that are released into lacteals of the villi eventually return from the systemic circulation to the liver where hepatocytes internalise the chylomicron remnants and recycle their components.^25^ We performed RNA-seq on liver samples from the same mice that we had previously performed ileal RNA-seq. We observed, similarly to the ileum, that broad transcriptomic profiles of MM12 mice were more similar to GF mice than to SPF mice (**Fig. 6a**). Consequently, relative to SPF, we found 1575 and 1963 genes significantly differentially expressed in GF and MM12 liver, respectively (**Fig. 6b-c** and **Supplemental Tables 8-9**). Significantly differentially expressed genes in both groups represented overlapping pathways (**Fig. 6d**). Given our previous analyses in ileum, and recent evidence for microbiome control over lipid uptake and transport^14^, it was notable that we observed a significant enrichment for genes involved in cellular responses to lipid (GO BP Response to lipid) among genes that were increased in both GF and MM12 liver samples (**Fig. 6d**) relative to SPF. Leading edge genes in both comparisons (GF vs. SPF and MM12 vs. SPF) represented canonical hepatocyte responses to chylomicron remnant uptake, including master transcription factors in lipid metabolism (*Cebpb* and *Ppargc1a*, **Fig. 6e-f**), as well as fatty acid uptake from chylomicron remnants (*Ldlr*, **Fig. 6g**). Relevant to previous literature, we also observed increased expression of *Cyp7b1* (**Fig. 6h**), the rate-limiting enzyme in the alternative bile acid synthesis pathway, supporting a role for increased chylomicron uptake in activating bile acid synthesis.

**Fig. 6.**
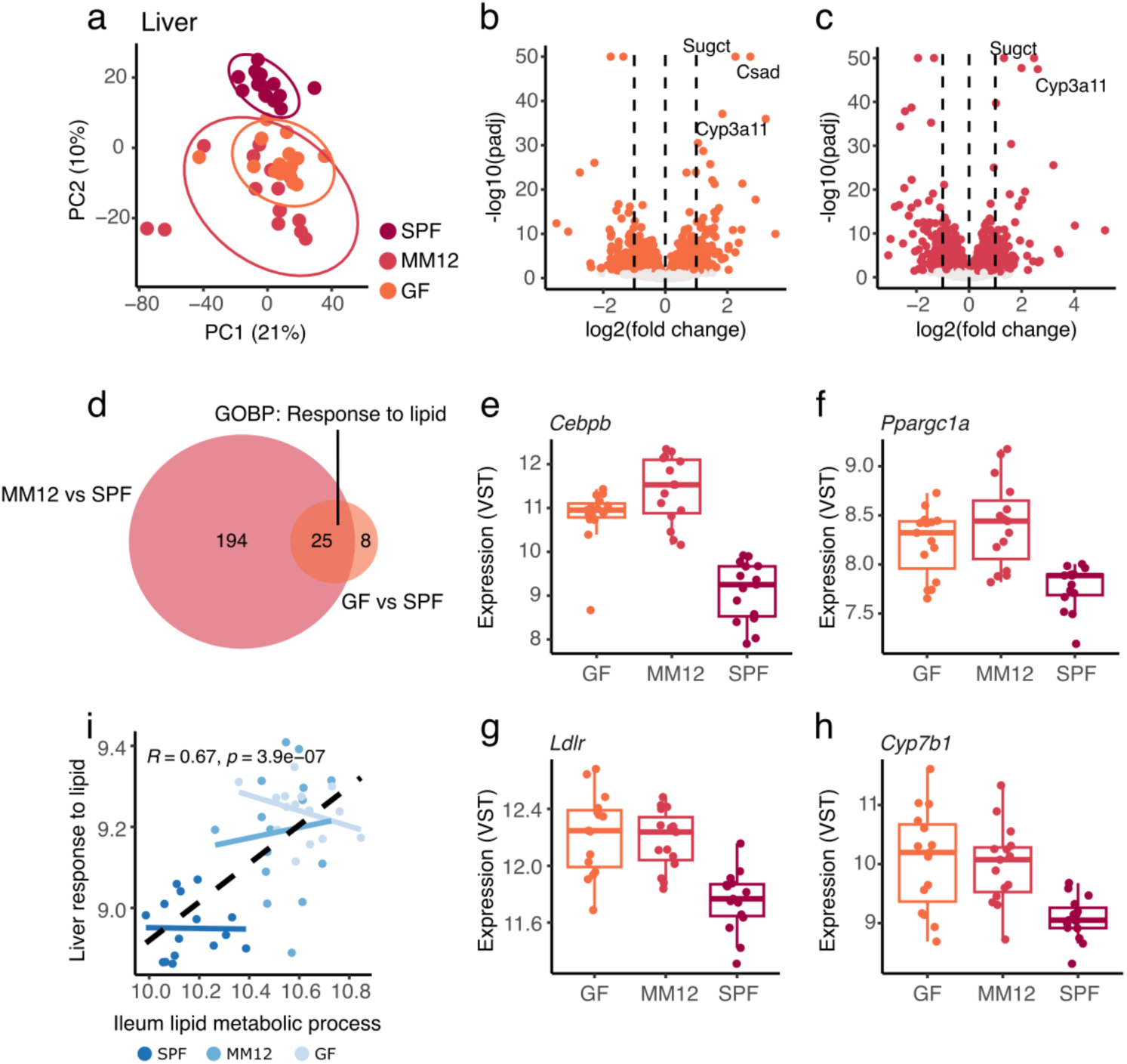
**Microbiome complexity impacts transcription of metabolic genes in the liver**. (a) PCA of liver transcriptomic profiles in GF, MM12 and SPF mice. Volcano plots showing differential gene expression results for (b) GF vs. SPF and (c) MM12 vs. SPF are shown. Coloured points = pad < 0.05 and guidelines at |log2 fold change| = 1 are shown. (d) Venn diagram showing the overlap of significantly enriched pathways in GF vs. SPF and MM12 vs. SPF comparisons (padj < 0.05). Representative genes annotated as being part of GO BP Response to lipid pathway are shown in (e)-(h) and represent master transcriptional regulators (*Cebpb* and *Ppargc1a*), lipid transport (*Ldlr*), and bile acid synthesis (*Cyp7b1*) genes. (i) Correlation between a metagene of genes annotated as GO BP lipid metabolic process in ileum and GO BP response to lipid in liver.

These data support a role for increased lipid uptake^14^ and induction of lipid metabolic pathways and chylomicron formation in GF and MM12 mice that may lead to an increased uptake of chylomicron remnants from the circulation. Indeed, we observed a significant group-shift in genes related to lipid metabolism in both GF and MM12 ileal samples with a concomitant increase in genes involved in responses to lipid in liver samples. However, the relationship was not observed within groups (**Fig. 6i**).

Together, these data support and extend recent evidence^14^ that the microbiome suppresses lipid absorption in the small intestine with a resultant change in liver metabolic transcriptional phenotypes.

## DISCUSSION

A large body of research has explored the role of the microbiome in mediating colonic immune function, digestion and nutrient absorption. However, due to challenges of access in human participants, knowledge of the small intestinal microbiome and therefore how it impacts mammalian health is less developed.^26^ However, segments along the small intestinal tract are critical for nutrient absorption and distribution, as well as feeding signals back to the liver. Therefore, and alongside regulation of immune system development and activation, the microbiome is a key player in regulating metabolic aspects of small intestinal function.

The complexity and composition of the small intestinal microbiome impact host processes. This is apparent from targeted studies that have shown that where GF mice have impaired secondary lymphoid structure development^10^ and increased lipid absorption^14^, these differences can be tuned by adding or subtracting specific species from the community, or through colonisation of minimally diverse communities. For example, using MM12 consortium, Brunner *et al.* were able to determine that the presence of a microbiome suppresses lipid absorption through a bile acid-dependent manner, yet some features of fatty acid and bile acid metabolism are intermediate between GF and SPF mice.^14^ This is in keeping with previous studies that have reported intermediate immune phenotypes in the colon and circadian dependence of MM12 metabolic phenotypes.^19^ Nevertheless, global impacts of small intestinal microbiome complexity on functional aspects of the host have not been explored.

Given this lack of systems-level information on the role of small intestinal microbiome complexity on host functions, we chose to transcriptionally profile the microbiome and host tissue in the terminal ileum (the major region of bile acid reuptake) across groups of mice with increasing levels of ileal microbiome complexity (GF, MM12 and SPF). We have demonstrated that while the MM12 community was primarily designed to recapitulate colonic microbiome taxonomy and function,^16^ it captures the broad taxonomy of SPF ileal microbiomes with a predominance of *Lactobacillus spp.* A major difference in MM12 relative to SPF animals is increased relative abundance of *Akkermansia muciniphila*, a species that is well-studied for its roles in mucin degradation, immunogenicity and metabolic properties, ^27–29^ and is a key bacterium in tuning host lipid metabolism.^14^

Assembly of microbial communities is determined by multiple factors that includes nutrient requirements of individual organisms and extent of metabolic competition.^30^ It is of note that competition between MM12 members *in vitro* is dominated by inhibitory effects and outgrowth of *Enterococcus faecalis* KB1.^31^ We do not see dominance of *E. faecalis* KB1 in the murine ileum, which suggests that the assembly dynamics, metabolic requirements, and metabolic outputs are crucially dependent on the *in vivo* environment. We attempted to assess this further using MSMM implemented using MICOM.^20^ We found that in contrast to the SPF community, the predicted metabolite production capacity of the ileal MM12 was limited. Enrichment of central metabolic functions in shared metabolites (i.e. pyruvate metabolism) suggested more limited secondary metabolism in MM12 mice. This may be explained by lower competition and more discrete niche occupancy in the minimally diverse MM12 community. Indeed, our analysis supported more consistent import metabolite niches across samples in the MM12 and more well-defined species-specific niche occupancy. Conceptually, the higher diversity and potential for niche clashes in the SPF community may require more diverse utilisation strategies and thus the perhaps inadvertent production of more metabolic intermediates.

The differences in complexity and metabolic outputs may explain some of the observed differences between the MM12 community and the SPF community in terms of ileal transcriptomic profiles. Of note, the MM12 community is unable to recover the majority of gene expression changes observed in GF mice relative to SPF mice. This suggests that multiple key species or functions are lacking in this community. This is particularly evident for certain lipid handling processes (e.g. expression of *Ldlr* and *Fabp1*) as well as innate and adaptive immune responses that includes the expression of *Igha*. While the MM12 community contains a representative group of strains, it does not fully capture the immunogenicity of a more diverse community. This means that while MM12 strains that includes *A. muciniphila* may mediate IgA responses,^18^ immunogenic species such as SFB^32^ are not present and may be required for more robust IgA production as would be observed in SPF mice.

A benefit of the current study is that we included sufficient numbers of mice to build gene co-expression networks. In SPF mice we were able to build robust networks that represent core ileal processes. It is interesting that alongside absolute differences in gene expression due to microbiome complexity, we observed changes to gene co-expression network topology. For example, we observed reduced connectivity in a gene module that contained genes involved in muscle cell differentiation (ILE3 tan module), that includes genes such as *Pdgfra*, *Syne1* and *Cdh2*. This likely reflects the existence of a smooth muscle regulatory network involved in peristalsis and movement of intestinal contents that is not recapitulated in MM12 mice. It is noteworthy that GF mice have significantly longer transit time relative to SPF mice,^33^ although it is not clear where MM12 sit in terms of this phenotype. Our data point to the potential lack of microbial functions in the MM12 community which may affect small intestinal transit. We also found a well-connected module involved in fatty acid metabolism and chylomicron assembly which contained genes such as *Apoc2*, *Apoc3*, *Apoa4* and *Fabp1* (ILE11, green module). Consistent with previous data,^14^ we find that the MM12 re-establishes connectivity that is lost in GF mice for this module. However, it is noteworthy that not all genes in this network are recapitulated in MM12, which speaks to nuances in microbiome complexity and composition which are incompletely understood at present. In our study, we have identified a candidate bacterium, *Faecalibacterium duncaniae*, which is associated with multiple host co-expression modules in the ileum. It is positively correlated with genes involved in fatty acid metabolism and glucose liberation and negatively correlated with genes upstream of innate immune responses, including bile acid signalling. These data fit well with previous literature that has uncovered the association between bile acid homeostasis and *Faecalibacterium spp.*.^34,35^ and it’s anti-inflammatory properties^35–37^. *Faecalibacterium spp.* are well-studied as potential probiotic species,^38^ with *F. duncaniae* (strains of which are re-classified *F. prausnitzii*) being associated with the production of protective metabolites^39^ and skewing towards IL10-production and regulatory response.^37,40–42^ Our data support pro-lipid and anti-inflammatory properties of *F. duncaniae* in the mouse ileum although the direction of effect or causal relationship have not been determined.

Finally, we have extended previous findings in microbiome regulation of the small intestine-liver axis. Transcriptomic data in mouse liver show increased expression of genes involved in response to lipid pathways and chylomicron remnant intake and handling (e.g. *Ldlr, Cebpb, Ppargc1a*) in GF and MM12 mice relative to SPF. This may be a consequence of increased lipid uptake and lymphatic/systemic transport in mice with no or low diversity microbiomes. Although MM12 can restrict lipid uptake to levels similar to SPF mice.^14^ this metabolic feature will, at least in part, be determined by the relative abundance of species which are capable of stimulating the upstream alternative hepatic bile acid synthesis pathway.^14^

### Limitations

The major limitations of the study are that we performed experiments in a single sex of animals. There are a number of studies that support sex-dependent effects in GF phenotypes, and our observations and interpretations should be extended to male animals with caution. MSMM is a powerful tool to explore predicted metabolic flux. However, there are limitations to interpretation of its outputs. We did not model the host in combination with the microbial communities and therefore could not explore the complex interactions between the intestinal epithelium and bacterial metabolites for example. A lack of host model may also inflate the predicted ability of bacteria to access and import metabolites from the environment. Our predictions of increased production of metabolites from central metabolic processes such as pyruvate metabolism require validation using mass-spectrometry or nuclear magnetic resonance-based methods. Co-expression network analysis has its challenges, especially when comparing strength of correlations between groups. We included the minimum recommended number of animals (n = 15) to provide robust estimates of correlation between genes. However, more samples would have provided more accurate results. While we could find some evidence of changes to correlation structure between groups of mice, we remain cautious in over-interpreting differences where sampling noise can influence such structure. Finally, we have provided evidence for an association between a well-studied bacterium, *F. duncaniae* with multiple host modules. The reasons for these associations should be interpreted with caution as we do not provide any causal links between bacteria and host. In addition to causal relationships, correlations could easily be due to factors (e.g. variation in bile acid composition) that independently impact on the relative abundance of *F. duncaniae* and the expression of ileal metabolism-related genes.

## METHODS

### Animals

All mouse (Mus musculus) work was conducted at the Biomedical Services Unit (BSU) within The Kennedy Institute of Rheumatology (KIR) at the University of Oxford. Mice were provided with sterile water and irradiated food ad libitum. Mice were culled via asphyxiation using a rising concentration of carbon dioxide, authorised via Schedule 1 of A(SP)A 1986, and confirmed via cervical dislocation.

Specific pathogen free (SPF, n = 15) mice were acquired from Charles River Laboratories (USA) and housed in the BSU at the KIR. Germ-free (GF, n = 15) mice were derived and maintained within the germ-free facility at the KIR and were kept in flexible film isolators under positive pressure. Oligo-mouse microbiota 12 (MM12, n = 15) mice were provided by Prof. Andrew Macpherson (Bern, Switzerland) and routinely screened for contamination through metagenomic sequencing. They were maintained within the germ-free facility and were kept in flexible film isolators, in the same manner as the GF mice. For all groups of mice, schedule 1 procedure was conducted at 12 weeks old. To reduce impact of cage effects in downstream analysis, multiple cages of animals were used per group. 3 (SPF and MM12) or 4 (GF) cages were used per group, housing 2-7 mice per cage. All sample collections were carried out in the afternoon between 1-3pm. All mice were female.

### DNA extraction from ileal contents

DNA and RNA were co-extracted from ileal contents samples using the AllPrep PowerFecal Pro DNA/RNA Kit (QIAGEN, Germany). Samples were thawed on ice and lysed in PowerBead Pro Tubes containing 650ul Solution CD1 and 100ul phenol-chloroform-isoamyl alcohol (25:24:1) (Sigma-Aldrich, USA). Mechanical lysis was performed using the TissueLyser II (QIAGEN, Germany) at 25Hz for 2×5min. Lysates were centrifuged, and supernatants were processed through sequential inhibitor removal and separation steps using CD2 and CD3 buffers.

DNA was captured on MB DNA Spin Columns, while RNA was purified from the flow-through after ethanol addition using MB RNA Spin Columns (QIAGEN, Germany). Both nucleic acid fractions were washed and eluted in RNase-free water (QIAGEN, Germany). DNA and RNA was kept on ice during processing and subsequently frozen at -70°C until assays were performed.

### Quantification of the MM12 community via qPCR

MM12 members were quantified using Quantitative PCR (qPCR), performed using Takyon qPCR Low ROX Probe 2X Master Mix (Eurogentec, Belgium). One MM12 sample was not available and so microbiome analysis was performed on 14 samples in total. Each 10ul reaction contained 5ul Takyon Master Mix, 0.15ul of forward and reverse primer mix (each at 40uM; Integrated DNA Technologies, USA), 0.25ul of probe (10uM; Integrated DNA Technologies, USA), 3.6ul of nuclease-free water (Qiagen, Germany), and 1ul of DNA template. Final concentrations were 300nM for each primer and 250 nM for the probe (a 3’ fluorescent quencher (TAMRA dye) conjugated with a 5’-labelled with 6-carboxyfluorescein (FAM)). Water volume was adjusted if higher DNA input was required. Primers and probes used in the reactions are provided in **Supplemental Table 10**.

Master mix (9\,µL) was dispensed and 1µL DNA was added. Plates (Thermo Fisher Scientific, USA) were sealed and briefly centrifuged to ensure even mixing. qPCR was run on a Viia7 system (Applied Biosystems, USA) using the following cycling conditions: 95°C for 3 minutes, followed by 45 cycles of 95°C for 15 seconds and 60°C for 1 minute. All reactions were performed in technical triplicates and included a standard curve on each plate. Sample and target annotations were applied during setup and output files were exported after completion for further analysis.

Standard curves were constructed from linearised plasmids harbouring 16S sequences of MM12 members. Plasmid maintenance involved transformation of Subcloning Efficiency DH5〈 Competent Cells (ThermoFisher Scientific, US) in SOC media (ThermoFisher Scientific, US) following manufacturer’s protocols. Post-transformation, cells were plated on LB agar containing 100ug/mL ampicillin and incubated overnight at 37°C. Individual transformed colonies were inoculated into LB broth supplemented with 100ug/mL ampicillin and cultured overnight at 37°C. Plasmid DNA was isolated using the Isolate II Plasmid Mini Kit (Meridian Bioscience, US) per manufacturer’s guidelines. Linearisation was achieved through restriction digestion using *NotI*, *HindIII*, or *NcoI* enzymes (New England, USA) as appropriate for each Oligo-MM12 member primer/probe combination, following manufacturer’s protocols. Plasmid standards were generated via 10-fold serial dilutions starting from 2 x 10^8^ copies/ul.

### Metagenomic sequencing of the SPF community

Metagenomic DNA quality control and library preparation were performed by Novogene (Beijing, China). DNA concentration and purity were assessed using the Agilent Fragment Analyzer 5400 (Agilent Technologies, USA). Libraries were prepared using a proprietary Novogene in-house kit, with an input of 100 ng DNA. Individual libraries were quantified using the Qubit Fluorometer (Thermo Fisher Scientific, USA), and size distribution was assessed using the Agilent 2100 Bioanalyzer (Agilent Technologies, USA). Final library concentrations were determined by qPCR. Sequencing was performed on the NovaSeq X Plus platform (Illumina, USA) using a paired-end 150 bp (PE150) strategy.

### RNA extraction from ileum and liver tissue

Terminal ileum and liver samples were removed and stored in RNAprotect (Qiagen, The Netherlands) at -70°C. Tissues were defrosted, weighed, and homogenised using a Precellys Lysing Kit – soft tissue ceramic beads (Precellys, France) in a TissueLyser II (Qiagen, The Netherlands) for 10 mins at 30Hz. RNA was extracted using the RNeasy Mini Kit (Qiagen, The Netherlands) as per manufacturer’s instructions. On-column DNase digestion and RNA clean-up was also conducted as per manufacturer’s instructions. Final RNA quantity and purity was checked via NanoDrop (Thermo Fisher Scientific, USA).

### RNA sequencing

RNA was plated and sent to the Oxford Genomics Centre (OGC, Centre for Human Genetics, Oxford) for library preparation and sequencing. Starting RNA was quantified using RiboGreen (Invitrogen, USA) on the FLUOstar OPTIMA plate reader (BMG Labtech, Germany) and the size profile and integrity analysed on the 2200 or 4200 TapeStation (Agilent, RNA ScreenTape, USA). RNA Integrity Number (RIN) was estimated to be between 5.2 and 10 for all samples, and RNA was normalised to 100 ng prior to library preparation. Polyadenylated transcript enrichment as well as strand-specific library preparation was performed using NEBNext Ultra II mRNA kit (NEB, USA) as per manufacturer’s instructions. Library amplification was done on a Tetrad (Bio-Rad, USA) for 14 cycles using a custom dual indexing primers based on previous literature.^43^ Qubit normalisation was conducted on individual libraries, and the size profile was analysed via TapeStation followed by pooling. The pooled library was diluted to approximately 10 nM for storage as well as denaturation and further dilutions prior to loading on the sequencer. Paired-end sequencing was performed using a NovaSeq6000 platform (Illumina, NovaSeq, USA) using the 6000 S2/S4 reagent kit v1 for 300 cycles (150bp paired-end).

### Metagenomic data processing

Preprocessing of shotgun sequencing data was performed using pipeline_preprocess from OCMS Shotgun (https://github.com/OxfordCMS/OCMS_Shotgun) which wraps functionality from multiple tools. Briefly, duplicates were removed using CD-HIT^44^ with default parameters, adapters were trimmed using Trimmomatic^45^ (parameters: 3 seed mismatches, 30 palindromic match score, 8 minimum adapter length, 25 quality prior to trimming, 36 minimum read length), reads were dusted using dustmasker^46^ with default parameters and low-complexity reads were masked with BMTagger^47^ with default parameters. Contaminating host reads were removed by aligning sequences to both human Hg38 and mouse Mm39 genomes using HISAT2.^48^

Taxonomic classification of filtered metagenomic reads was performed using Kraken2^49^ using the k2 pluspf 20230605 database with –confidence = 0.7. Counts at each taxonomic level were generated using Bracken^50^ (parameters: read length 150). Selection of the confidence parameter for Kraken2 was conducted using two ZymoMock community standards (https://www.zymoresearch.com/collections/zymobiomics-microbial-community-standards/products/zymobiomics-microbial-community-standard) where results were compared to the expected results (**Supplementary Material Fig. S3**). Taxa with a relative abundance of over 0.001% in more than 3 samples were included in downstream analyses. Principal Co-ordinate Analysis (PCoA) was conducted using the ordinate() function from phyloseq v1.44^51^, using Bray-Curtis distance. Shannon diversity was calculated using estimate_richness() from phyloseq v1.44.

### Metagenome-scale metabolic modelling (MSMM)

Genome-scale models for SPF and MM12 communities (strain level) were curated from the AGORA2 database.^52^ In cases where models for the exact strains were unavailable, the ATCC or DSM strain were used as representatives. If no genome-scale models were available for the species, genomes and genome annotations were sourced from RefSeq,^53^ and were constructed with CARVEME v1.6.5.^54^ Reconstruction used the bacteria universal model, with protein sequences supplied in FASTA format and scored by DIAMOND against the BiGG reaction database. Draft models were gap-filled, such that each model was required to carry flux through its biomass reaction on that medium. All other parameters were left at package defaults. Because AGORA2 models are built in the VMH namespace and CarveMe models in BiGG, exchange reaction identifiers in the CarveMe models were mapped to VMH identifiers prior to community assembly. We reconstructed models for *Duncaniella sp* C9, *Duncaniella sp* B8, *Duncaniella freteri, Duncaniella dubosii*, *Candidatus savagella* (SFB), *Sporofaciens musculi, Romboutsia ilealis, Rhodococcus fascians, Muiribaculum gordoncarteri, Moraxella osloensis*, *Ligaliactobacillus murinus, Lactobacillus sp* AMBV1719, *Lactobacillus paragasseri*, *Faecalibaculum rodentium* and *Faecalibacterium duncaniae*.

Mice were fed a standard mouse chow diet, SDS RM3 (pellet form) (LBS Biotechnologies, United Kingdom). Using the product information sheet, the diet was converted into a format appropriate for metabolic modelling. The composition (g/kg feed) was converted into a MICOM medium file: exchange reaction IDs paired with maximum uptake flux in mmol gDW⁻¹ h⁻¹, derived by converting mass to moles, scaling to 4.5 g/day intake and dividing by 24. We did not measure the consumption of food, thus we estimated a daily consumption of 4.5g and calculated from that the daily consumption of each nutritional component. Host-supplied components such as mucins were added as per the MICOM diet repository, as well as appropriate dilutions applied to the nutritional component to simulate absorption.^20^ Minerals, trace elements and water-soluble vitamins were scaled by 0.5 per the general mouse metabolic model and the ; host-supplied and poorly absorbed components (oxygen, glycine, core oligosaccharide, conjugated bile acids) were undiluted; all others retained diet-derived flux. Additionally, we used a general murine metabolic model to simulate the production of host components such as bile acids and to estimate the consumption of metabolites prior to ileal metabolism.^55^ This resulted in the creation of a custom mouse-specific dietary input for metabolic modelling that attempted to recapitulate the metabolic environment available to the microbiome in the ileum (**Supplemental Table 11**).

Metagenome-scale metabolic modelling was conducted using MICOM v0.37.1.^20^ We determined the cooperative tradeoff value - the balance between optimising for community versus individual taxa growth - through the inbuilt tradeoff() function. For both SPF and MM12 communities, we selected the value that allowed for significant growth of all bacteria in the community. This was set at 0.5 and applied uniformly across all samples within each cohort. We then used the inbuilt grow() workflow function to simulate microbial growth on the diet. Total fluxes were exported for downstream analysis. Analyses were performed using iPython notebooks using Python v3.10. Gurobi 10.0.1 (GCCcore 12.2.0) was used as the optimisation solver where applicable. We used the predicted export fluxes for downstream analyses, representing bacterial biosynthetic production. For each metabolite in each sample, bacterial production capacity was the sum of export-direction exchange fluxes across all taxa, representing gross output before intra-community consumption. Net community-environment exchange was the community-level (medium) flux. Both were filtered at ≥3 samples and ≥0.0001 mmol gDW⁻¹ h⁻¹.

Pathway enrichment analysis for overlapping and distinct predicted metabolites between SPF and MM12 communities was carried out using the KEGG compound and pathway databases^56^ and overrepresentation analysis implemented using the fora() function in fgsea v1.28.0.^57^

### RNA-seq data processing and analysis

FastQC v0.12.0 and MultiQC v1.14^58^ were used within the Oxford Centre for Microbiome Studies (OCMS) RNA-seq pipeline to run quality control on the raw FastQ files. Pseduoalignment against the mouse transcriptome (genome assembly GRCm39 mm39, ENSEMBL build 107) was performed using Kallisto v0.46^59^. Tximport v1.28.0^60^ was used to collate the gene counts from all samples. Quantification at gene-level was used for all downstream analyses. Genes were filtered out if they had fewer than 10 reads in fewer than 15 samples. DESeq2 v1.40.2^61^ was used to conduct variance stabilising transformation (VST) on the read counts and transformed data were used for exploratory analysis and visualisation.

Principal component analysis (PCA) was conducted using the prcomp() function in base R v4.3.2 to look for structure in the data. Analysis of ileal data revealed major contributions to variation in gene expression that was not attributable to microbiome status but to some inter-sample variation (**Supplementary Material Fig S4a**). To assess the reasons for this structure, we added ileum transcriptomic data from an extension of the current project (an aging analysis) to include a total of 90 ileum samples that were representative of different groups of animals (**Supplementary Material Fig S4a**). While some of the variation may have been attributed to RNA quality (**Supplementary Material Fig S4b**), we observed that genes with the highest PC1 loadings were genes involved in lipid and nutrient handling (*Lct, Fabp1, Apoa4*, **Supplementary Material Fig S4c-e**). This suggested that the main sources of variation were due to sampling variation through subtle tissue differences or factors such as time-since-feeding, which we did not measure. To remove this unwanted source of variation, we removed the variation attributed to PC1 and PC2 using limma^62^ removeBatchEffects() for visualisation purposes (**Supplementary Material Fig S4f**) and included them as covariates in downstream differential expression analysis.

DESeq2 v1.40.2^61^ was used for differential expression between groups (pairwise tests of GF vs. SPF, MM12 vs. SPF and GF vs. MM12). Benjamini-Hochberg adjusted p-value < 0.05 was used to define significantly differentially expressed genes. Gene set enrichment analysis (GSEA) was conducted using the fgsea v 1.28.0 (fgsea function) using annotations from Gene Ontology:Biological Pathways (GOBP) pathway database (m5.go.v2023.1.Mm.symbols.gmt.txt from mouse MSigDB). Significantly enriched pathways were defined as those with a Benjamini-Hochberg adjusted p-value < 0.05.

### Weighted Gene Co-Expression Network Analysis (WGCNA)

Weighted gene co-expression network analysis (WGCNA) was conducted using the R package WGCNA v1.73^65^ where we constructed signed networks in data from ileal samples of SPF mice. Input into WGCNA was VST-transformed data. Soft-power thresholds were determined by assessing scale-free topology and mean connectivity, resulting in the use of the soft power threshold of 7 for the ileum. Resulting WGCNA-generated co-expression modules were merged at 80% co-expression similarity. The modules were annotated using enrichment analysis of GOBP in clusterProfiler^63^. Module preservation between groups was assessed using the modulePreservation() function in WGCNA, using signed network type for 500 iterations to get the final Z-score summary, hub stability, and co-expression scores. Correlations between species relative abundance and host modules was performed using Spearman rank correlations using species relative abundance and module eigengene values as inputs.

### Data availability

Raw RNA-seq data are available at ArrayExpress hosted at EMBL-EBI with accession number E-MTAB-17749. Raw shotgun metagenomics data are available at the EBI ENA with accession number PRJEB127927/ERP207389. Code used to analyse the data are available on github (https://github.com/OxfordCMS/Diversity_Transcriptome).

### Supplemental Tables

Supplemental tables are available at figshare (https://doi.org/10.6084/m9.figshare.35004151)

---

**Fig. S1.**
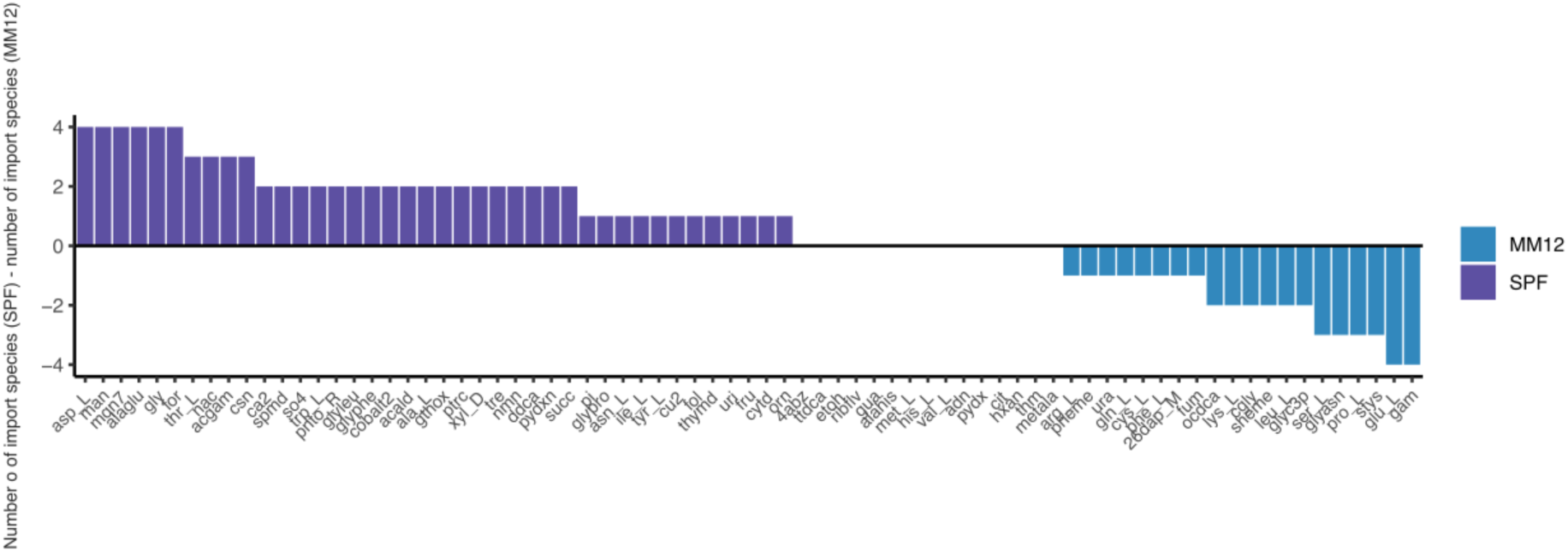
Metabolites are predicted to be imported by more SPF species than MM12 species. Y-axis represents the difference in the number of species predicted to import the given metabolite between SPF and MM12. Metabolites shown are those that are predicted to be imported at >0.00001mmol/gDW/h in at least 3 SPF and at least 3 MM12 samples.

**Fig. S2.**
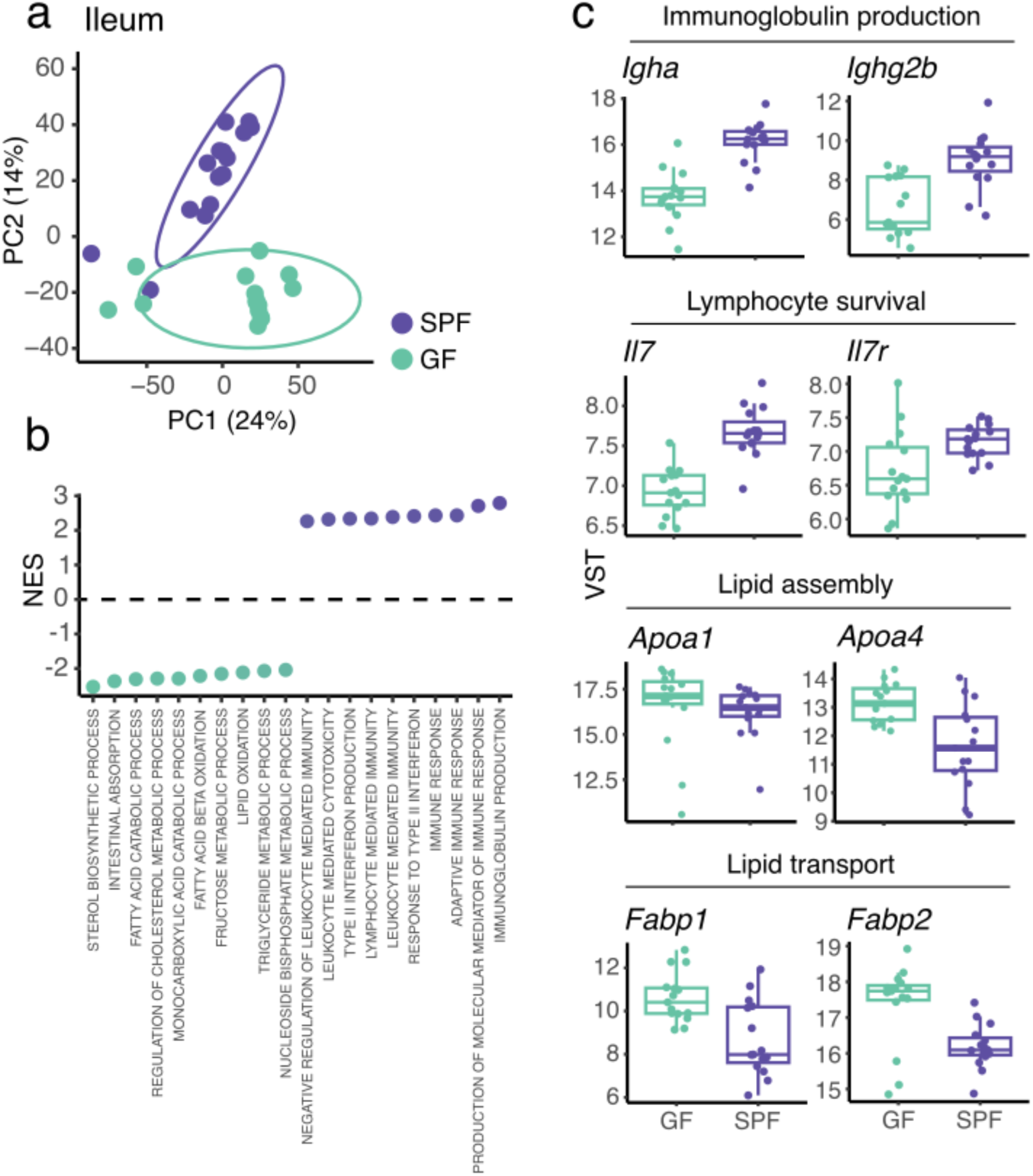
Ileal gene expression profiles in GF mice are different from SPF mice. **(a)** Principal components analysis (PCA) reveals global changes in transcription (based on VST expression values of 56,690 genes). **(b)** Top 10 significantly enriched pathways (padj < 0.05) from gene set enrichment analysis (GSEA) against Gene Ontology Biological Processes (GO BP) using genes ranked by fold change (14, 864 genes analysed in differential expression analysis). **(c)** Representative genes from up- and down-regulated gene sets which belong to the leading edge subset in top enriched pathways.

**Fig. S3.**
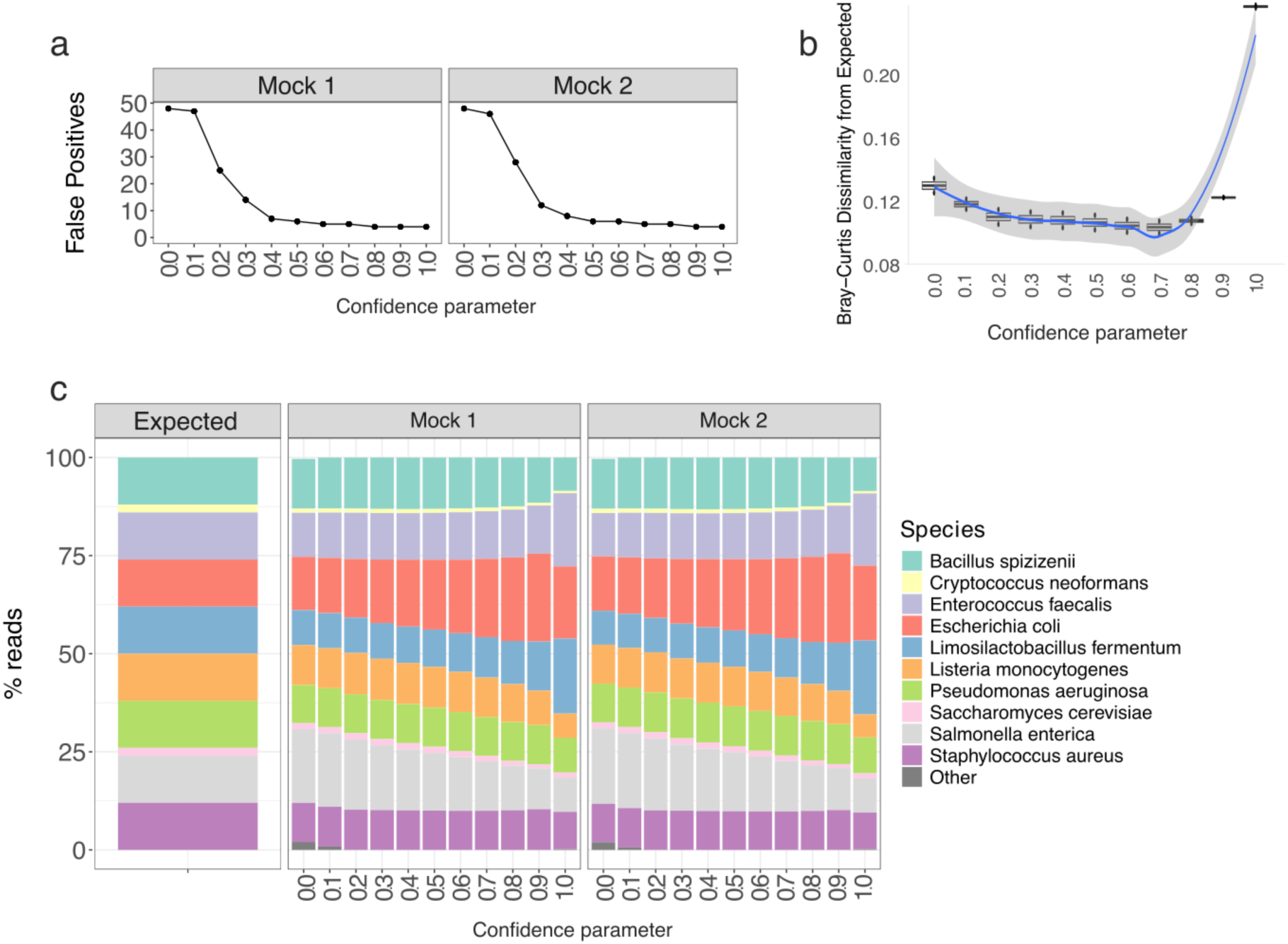
Shotgun metagenomic positive controls and kraken2 confidence parameter choice. We used two Zymo Mock communities to assess our ability to successfully extract DNA and assign species using kraken2/bracken. (a) Number of false positive species identified in two mock communities using varied confidence parameters in kraken2 from 0-1.0 (confidence parameter of 1 would require all annotations for a read to be the same species in order for it to be called as that species). (b) Bray-curtis dissimilarity of each mock sample at each confidence parameter. Dissimilarity decreases until confidence = 0.8. (c) Stacked barchart to visually show the difference between predicted community composition of each mock community standard to the expected (i.e uniform distribution).

**Fig. S4.**
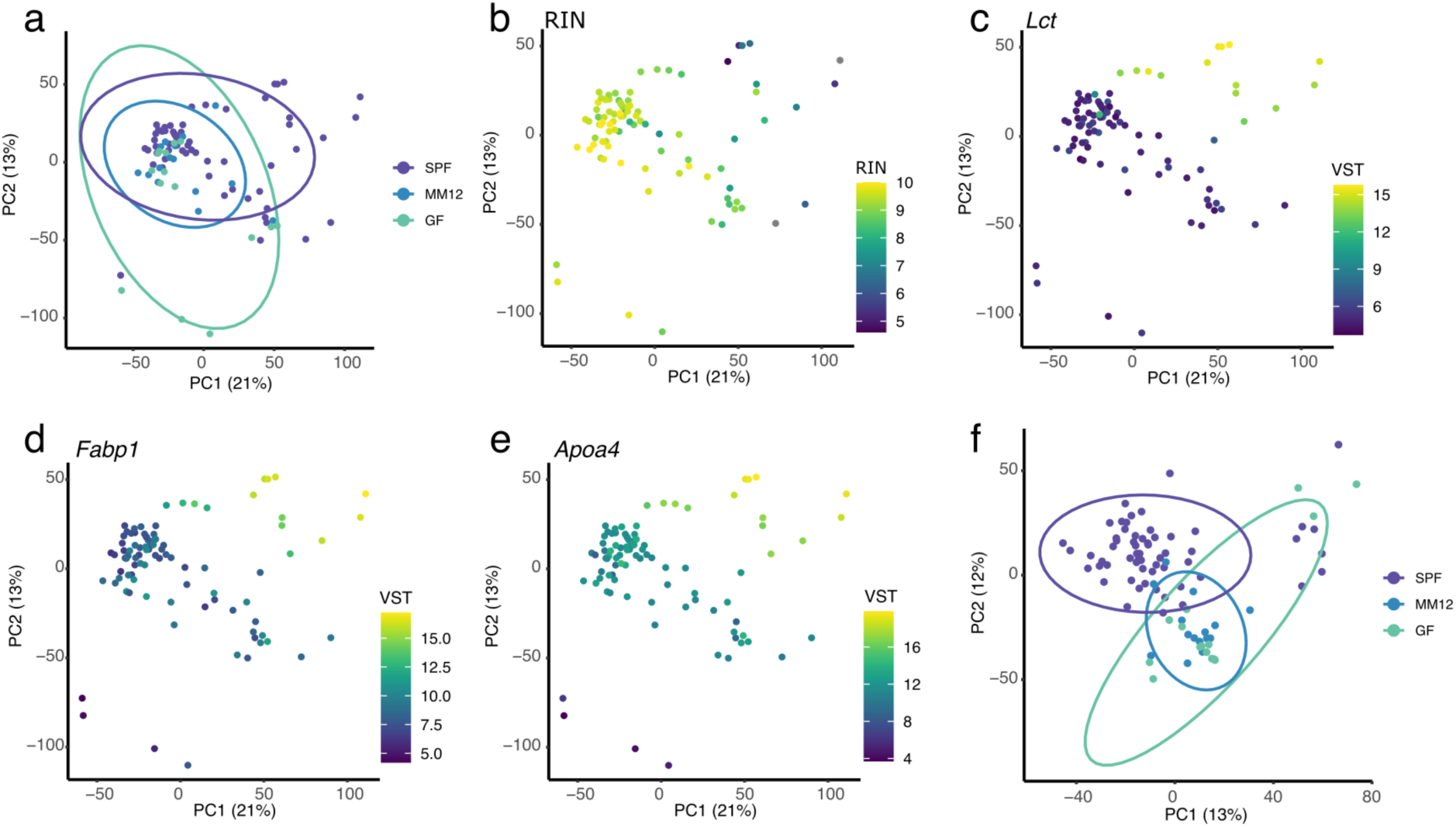
Correcting for interindividual variation in ileal gene expression. (a) PCA of ileum gene expression (VST) from 90 samples across multiple groups as part of a larger dataset than is presented in this manuscript. The data also include old (52 week), young (12 week) and old + young cohoused C57BL/6 mice which were sequenced alongside the reported samples. These additional datasets are available alongside the data presented in this manuscript at the EBI ENA with accession number E-MTAB-17749. PCA plots that with points coloured by (b) RNA integrity number (RIN), (c) *Lct* expression, (d) *Fabp1* expression and (e) *Apoa4* expression. (f) PCA of corrected gene expression (VST), corrected using the removeBatchEffects() function from limma v3.58.1 in R4.3.1.

